# Effects of oscillation phase on discrimination performance in a visual tilt illusion

**DOI:** 10.1101/2023.12.20.572703

**Authors:** Jessica G. Williams, William J. Harrison, Henry A. Beale, Jason B. Mattingley, Anthony M. Harris

**Affiliations:** School of Psychology, The University of Queensland, St Lucia, Australia; Queensland Brain Institute, The University of Queensland, St Lucia, Australia; Canadian Institute for Advanced Research (CIFAR), Toronto, Canada

**Author notes:** Correspondence should be addressed to Anthony M. Harris.

**Keywords:** Neural oscillations, Alpha oscillations, Tilt Illusion, Phase, Visual processing, Lateral inhibition, Excitation/Inhibition balance

## Abstract

Neural oscillations reflect fluctuations in the relative excitation/inhibition of neural systems^1–5^ and are theorised to play a critical role in several canonical neural computations^6–9^ and cognitive processes^10–14^. These theories have been supported by findings that detection of visual stimuli fluctuates with the phase of oscillations at the time of stimulus onset^15–23^. However, null results have emerged in studies seeking to demonstrate these effects in visual discrimination tasks^24–27^, raising questions about the generalisability of these phenomena to wider neural processes. Recently, we suggested that methodological limitations may mask effects of oscillation phase in higher-level sensory processing^28^. Thus, to test the generality of phasic influences requires a task that requires stimulus discrimination but depends on early sensory processing. Here, we examined the influence of oscillation phase in the visual tilt illusion, in which an oriented centre grating is perceived titled away from the orientation of a surround grating^29^. This illusion is produced by lateral inhibitory interactions in early visual processing^30–32^. We presented centre gratings at participants’ titrated subjective vertical angle and had participants report whether the grating appeared tilted leftward or rightward of vertical on each trial while measuring their brain activity with EEG. We observed a robust fluctuation in orientation perception across different phases of posterior alpha and theta oscillations, consistent with fluctuating illusion magnitude across the oscillatory cycle. These results confirm that oscillation phase affects complex processing involved in stimulus discrimination, consistent with their purported role in canonical computations that underpin cognition.

## Results

To probe early visual processing for potential influences of neural oscillation phase in tasks more complex than simple detection, we exploited an orientation-contingent visual illusion called the direct tilt illusion^29,33,34^ (hereafter, the *tilt illusion*; **Figure 1A**). The tilt illusion is a well-studied phenomenon in which a central test stimulus has its perceived orientation biased away from the orientation of a surround stimulus (i.e., a repulsive effect) when the angular difference of the surround relative to the centre is in the range of 10-40°. The tilt illusion is produced by suppressive lateral interactions in early visual cortex, whereby neural responses to a central test grating are suppressed in an orientation-specific manner by the presence of an oriented surround^30–32^. The orientation-specific nature of this effect suppresses population neural responses representing angles similar to that of the surround, producing a population response to the test stimulus that has its overall orientation distribution biased away from the angle of the surround^34^. The tilt illusion is an ideal candidate for probing modulation by neural oscillations because it is robust over short presentation times^35^, allowing for presentation during discrete portions of the oscillatory cycle.

**Figure 1.**
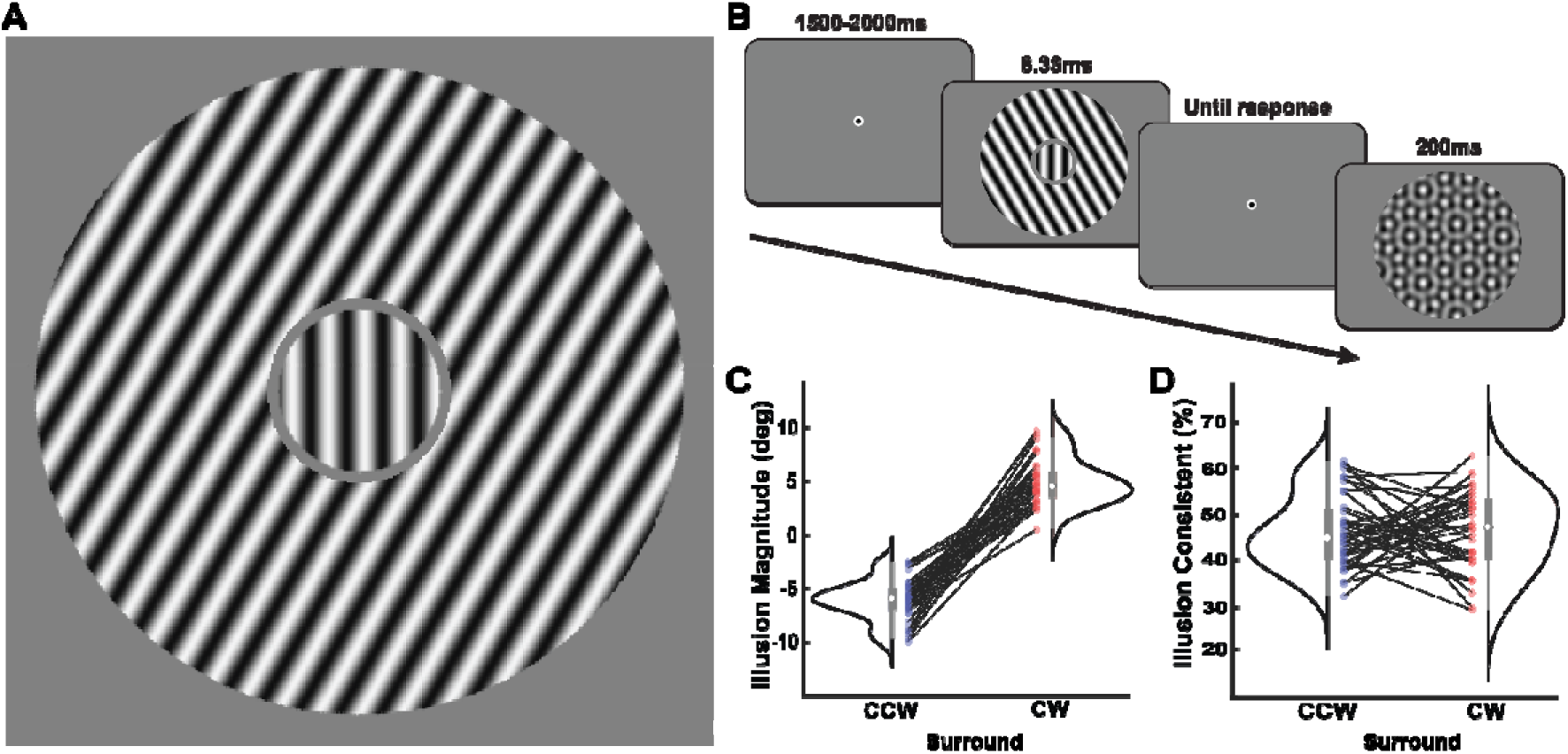
Task design and behavioural results. **A.** A demonstration of the illusion. The centre grating is oriented vertically but is perceived as tilted leftward due to the influence of the oriented surround. **B.** Participants judged the orientation of a central grating, titrated to their subjective perception of vertical under the tilt illusion. Surround gratings were presented at ±30°. Following each response, a mask was presented to prevent the buildup of orientation aftereffects across trials. **C.** Density plots showing illusion magnitudes in each surround condition. **D.** The percentage of illusion-consistent responses in each surround condition. As the tilt illusion is a repulsive illusion, illusion-consistent responses were coded as responses in the opposite direction to the surround. In C and D, coloured dots and thin black lines represent individual participants. White dots represent medians, thick grey bars represent the interquartile range, and thin grey bars represent 2.5x the interquartile range. CCW: counterclockwise. CW: clockwise.

Figure 1B shows a schematic of the trial design. Gratings oriented ±30° were presented surrounding central gratings that had their orientation titrated to the threshold angle for a vertical percept (the angle at which the central grating was seen as rotated counter-clockwise [CCW] and clockwise [CW] equally often for a given surround). Stimuli were presented briefly (8.33ms) at 60% contrast and were clearly visible. Participants responded to the perceived orientation of the central grating on each trial (CCW or CW relative to vertical). For all analyses, responses from the two surround conditions were combined by recoding them relative to the repulsive nature of the illusion. Responses were coded as either ‘illusion consistent’ (i.e., CW responses when the surround was CCW, and vice versa) or ‘illusion inconsistent’ (i.e., CW when the surround was CW, and vice versa).

### Behavioural confirmation of the tilt illusion

We first quantified the magnitude of the illusion for each individual. Figure 1C shows the bias-corrected illusion magnitudes for each participant from the CCW- and CW-surround conditions. Both surround conditions produced an illusion in the expected direction, and the effect was comparable to the magnitude of tilt effects reported in past research research^35^ (CCW surround: *M (SD)* = 6.05° (1.86°), *t*(35) = 19.51, *p* < .001, 95% CI [5.42°, 6.68°]; CW surround: -4.90° (2.11°), *t*(35) = -13.93, *p* < .001, [-5.61°, -4.18°]). Uncorrected thresholds for each participant and for each condition are shown in **Table S1**. Change in threshold across the experiment was examined to assess the stability of the illusion across time. Thresholds reduced across the course of the experiment (CCW surround: *M_reduction_ (SD)* = 0.61° (1.33°), *t*(35) = -2.74, *p* = .010, [-1.06°, -0.16°]; CW surround: 0.39° (1.44°), *t*(35) = 1.98, *p* = .056, [-0.01°, 1.02°]), suggesting a degree of habituation over time. The proportion of illusion-consistent responses in each condition departed slightly from 50% overall (Figure 1D; CCW surround: 0.46 (.08), *t*(35) = -2.70, *p* = .011, [.44, .49]; CW surround: .47 (.09), *t*(35) = -2.16, *p* = .037, [.44, .50]), but produced comparable numbers of illusion consistent and illusion inconsistent responses in each condition, allowing us to conduct a well-powered analysis of the influence of phase on behaviour^36^.

### Influence of trial-wise excitation and inhibition

Previous work has established that individuals with greater suppression of V1 Blood-Oxygen Level Dependent (BOLD) responses to orthogonal orientations relative to aligned orientations show a larger tilt illusion^37^, possibly due to the role of neural inhibition in modulating the precision of orientation tuning in early visual cortex^38–41^. Here we extend these findings by demonstrating that the magnitude of the tilt illusion changes with trial-by-trial fluctuations in the relative level of neural excitation and inhibition. We examined three correlates of excitation/inhibition, namely, alpha power, and the slope and offset of the power-law function of the EEG power spectrum.

The amplitude of pre-stimulus alpha oscillations is a well-established correlate of neural inhibition^13,42–44^. For example, higher alpha power is associated with reduced perceptual indices of excitability^45^, reduced neural firing^1^, and reduced BOLD response^46^. Alpha power in area V1 also correlates with the strength of surround suppression evoked by stimuli of different orientations^47^. The parameters of the power-law function of the electrophysiological power spectrum, which is associated with aperiodic brain activity, also reflect the relative excitation/inhibition of neural systems^3,48^. These parameters correlate with several indices of neural excitation and inhibition^3^ including neural firing rates^49,50^ and BOLD signals^51^, and have recently emerged as robust correlates of processing in a number of cognitive domains^52–60^.

Here, we used the specparam toolbox^48^ to measure, on each trial, alpha amplitude, aperiodic slope, and offset in the 500ms prior to stimulus onset, averaged across a central-posterior region of interest (ROI; see Method and **Figure S1**). We used logistic mixed-effects modelling to estimate whether each of these parameters predicted participants’ perception of the central grating on single trials. We constructed a model with fixed effects of alpha power, aperiodic slope and aperiodic offset and a random effect of participants, with the goal of predicting single-trial perceptual reports (illusion-consistent or illusion-inconsistent perceived tilt). We observed significant prediction of trialwise responses from both alpha power (β = .05, *p* < .001) and aperiodic slope (β = .07, *p* = .046), both of which suggest that trials with higher levels of inhibition were more likely to result in illusion-consistent responses, consistent with an increased illusion magnitude on those trials. Aperiodic offset was not a significant predictor of participants’ tilt judgements (β = -.04, *p* = .244). An exploratory logistic mixed effects analysis performed at each electrode and corrected for inflation of the false discovery rate^61^ showed similar results: Aperiodic slope was a significant positive predictor of illusion-consistent responses at 3 posterior channels. Offset was a significant negative predictor at one posterior channel, and alpha power was a significant positive predictor of illusion-consistent responses over most of the scalp (58 out of 64 channels; **Figure S2**).

### Tilt illusion strength fluctuates with prestimulus phase

Having established that the magnitude of the tilt illusion varies with levels of relative excitation/inhibition from trial to trial, we next asked whether the effect also varies with changes in the relative level of excitation and inhibition within a trial, due to fluctuations in neural oscillations in the visual system. We employed the *phase opposition sum*^36,62^, a well validated and sensitive metric of oscillatory phase effects. The POS quantifies, at each time and frequency in the prestimulus window, the degree to which two different response categories (here, illusion-consistent and illusion-inconsistent responses), cluster at different phases of an oscillatory cycle. This metric showed that the likelihood of participants making an illusion-consistent or illusion-inconsistent response fluctuated with the phase of prestimulus alpha and delta-theta oscillations at our posterior ROI (Figure 2A). After correcting for multiple comparisons, we observed significant phase dependence for signals between 11 and 15 Hz from -520ms until -477ms, at 6 Hz from -391ms until -359ms, and at 3 Hz from -367ms until -324ms. Exploratory POS analyses across all electrodes yielded similar results (**Figure S3**).

**Figure 2.**
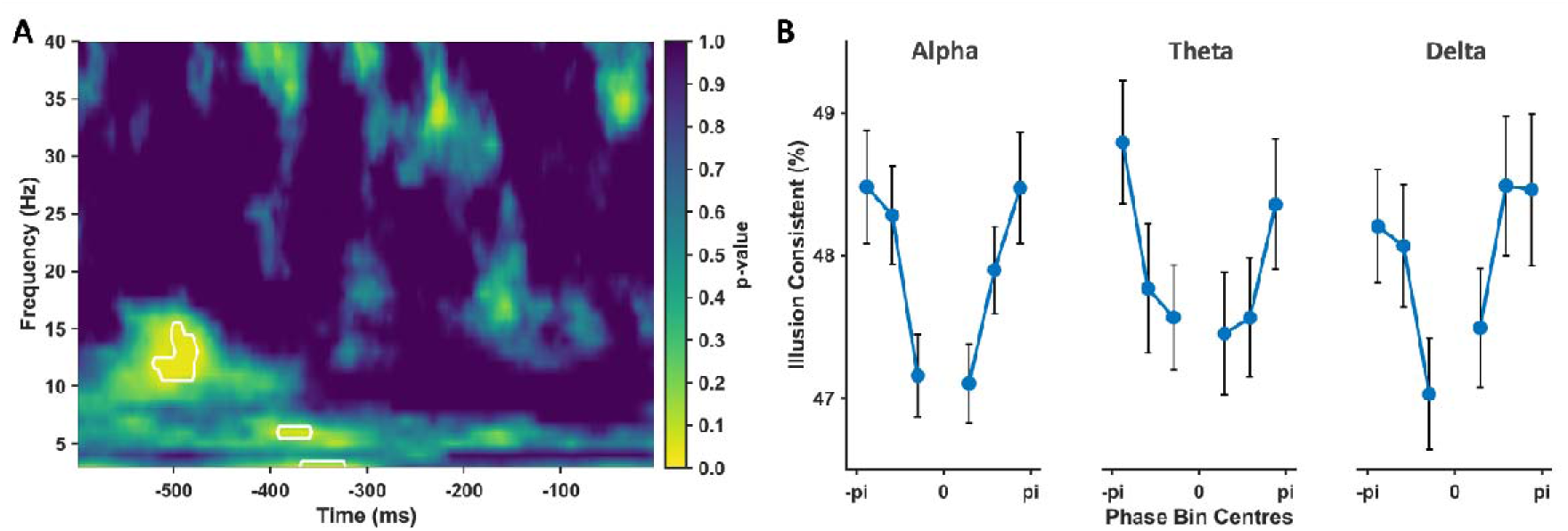
Phase opposition analysis shows tilt illusion magnitude depends on oscillation phase. **A.** Results of the phase opposition sum (POS) analysis in the pre-stimulus window (stimulus onset was at 0ms). Significant POS (indicated by white outlines) was observed in the alpha, theta, and delta range, after FDR correction for multiple comparisons^61^. **B.** Results of the unbiased phase alignment procedure, demonstrating the effects of phase on orientation perception in the tilt illusion. For each significant time and frequency, each individual’s responses were quantified for each of 7 phase bins of equal width. The phase bin with the fewest illusion-consistent responses was aligned to 0 radians and excluded from analysis. The remaining phase bins show significant modulation of tilt judgements by phase. Error bars represent within-participants standard error^63,64^.

We performed an unbiased alignment procedure to quantify the POS effects^15,17,20^. When results were aligned to the phase bin with the lowest proportion of illusion-consistent responses, all significant clusters showed the expected monotonic increase in illusion-consistent responses as phase deviated further from the aligned bin (Figure 2B). Comparing the bins nearest to the aligned bin with the most distant bins provided further evidence for modulation by phase in each case (alpha: *t*(35) = 2.63, *p* = .006; theta: *t*(35) = 1.81, *p* = .040; delta: *t*(35) = 1.83, *p* = .038).

### Influence of oscillatory power on detectability of phasic sampling

The analysis of phase effects in the prestimulus period is performed as a proxy for the effect of phase at the time the stimulus is processed. This approach allows us to avoid the confounding influence that stimulus-evoked responses have on phase estimates. An assumption of this method is that the neural oscillations relating to behaviour continue until the time the stimulus is processed. The effect of phase observed was relatively distant in time from stimulus onset (∼0.5 seconds), but we did not see a continuation of this effect until the time of stimulus onset. As such, we considered whether changes in oscillatory power may be masking the presence of phasic influences on behaviour in the intervening period. At very low power, oscillatory phase values become meaningless because the oscillation is effectively absent. However, even before this point is reached, when a low-power oscillation is present and may still be having physiologically relevant effects, phase values can become difficult to estimate as they are corrupted by the low signal-to-noise ratio of the data^65–67^.

Consistent with this proposal, we observed a significant decrease in the power of alpha oscillations over the prestimulus period, *t*(35) = 6.10, *p* < .001 (Figure 3), potentially explaining why phase effects were not observed throughout the prestimulus period.

**Figure 3.**
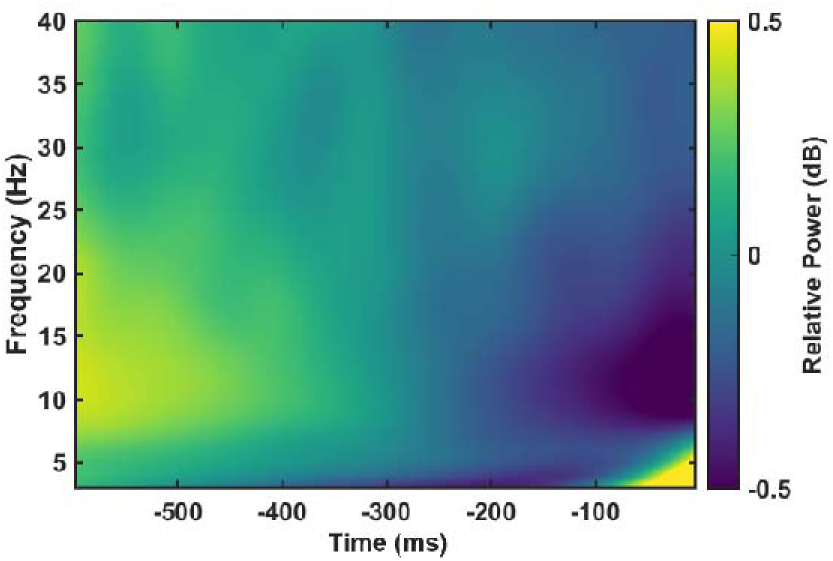
Change in power across the prestimulus window. Relative to the average of the prestimulus period, power in the alpha band (8-14 Hz) was higher at the start of the window relative to times closer to stimulus onset at 0ms.

## Discussion

We set out to test whether the phase of neural oscillations influences the inhibitory interactions in early visual cortex that give rise to the tilt illusion. We observed robust fluctuations in the likelihood that participants reported the central target as being tilted in a direction consistent with the illusion across the alpha, theta, and delta cycles. Our logistic regression results indicated the direction of these effects, with increased increased alpha power and increased aperiodic slope, both correlates of increased inhibition, corresponding to increased magnitude of the tilt illusion. The role of inhibition in producing orientation tuning in early visual areas is well established, although at least two different mechanisms for doing so seem to exist between, and potentially within, different mammalian species^31,38,41,68^. The current experimental design does not allow us to disentangle the mechanisms underlying fluctuations in tilt illusion magnitude. These fluctuations may be caused by fluctuations in the strength of the suppressive component of surround suppression across the oscillatory cycle. Alternatively, illusion magnitude may fluctuate if oscillation phase is influencing the precision of orientation tuning, modulating the degree of overlap in the population neural responses to different orientations. These possibilities are not mutually exclusive^69^.

Critically, our purpose in examining phasic influences on the tilt illusion was to test the involvement of neural oscillations in perceptual discrimination. We asked this question in response to recent null results in studies of phasic influences on perception^24,26,70,71^ that are difficult to explain given accounts of neural oscillations subserving canonical neural processes^6–8^. If phasic fluctuations in processing are fundamental to cortical computation, why are we not always able to observe their influence on behaviour? We have recently suggested that stimulus evoked brain responses, particularly the stimulus evoked phase reset, may place a fundamental limit on our ability to observe phase-behaviour relationships for anything but the most hierarchically early neural processes^28^ (see also^72^). To test this possibility we required a task in which the relevant neural processes occur early in the processing hierarchy but involve a level of complexity beyond the simple first-order detection processes linked to phase previously^15–17,20^. The tilt illusion is known to be caused by neural interactions in early visual processing^30–32^, making it an ideal candidate. Using this illusion, we were able to demonstrate that effects of phase on visual perception occur for high contrast, highly visible stimuli in a discrimination paradigm.

The influence of oscillation phase in discrimination performance is consistent with suggestions that neural oscillations play a role in processing throughout the cortical hierarchy^6^. Prior demonstrations of phasic influences on perception have primarily been in detection paradigms^15–17,20^. Detection paradigms test the simplest elements of sensory processing, requiring only that a threshold be placed on the activation of a neural population, which, if crossed, signals the presence of a stimulus. Whether the threshold is crossed due to a modulation of stimulus information or noise, or whether it contains any feature-specific content at all, is not tested by examining the effect of phase on hit rates^73^. Discrimination tasks, by contrast, are more complex, requiring comparison of feature-specific signal representations to multiple target categories, and evidence for the different categories to be accumulated. Demonstration of phasic influences on discrimination judgements shows an influence of neural oscillations beyond modulating general levels of activation in visual cortex. Our results demonstrate an influence of oscillation phase on interactions between disparate neural populations, consistent with suggestions of neural oscillation phase being involved in coordinating neural processing and communication^10–12,74,75^.

Recent null results in phasic sampling studies have also posed a challenge for oscillatory theories of neural processing and cognition. The power of alpha and theta oscillations has been linked to attention and perception in a wide range of different task contexts ^18,73,76–86^. Based on results such as these, and others showing neural oscillations influence neural signal transmission^10,11,87–90^, several theories have proposed that neural oscillations, and in particular, phase, play a key role in organising the neural processing that underpins cognition^11,12,14,42,74,91–94^. It is difficult for theories of rhythmic influences on cognitively-relevant neural processing to reconcile why such influences might not be observable in the outcomes of studies that probe those cognitive and perceptual functions. However, null results do not unequivocally contradict theory if those nulls may be due to methodological limitations. Our results point towards some key methodological limitations to be considered in studies of phasic influences on perception and cognition.

There are several ways in which an existing phasic influence may be invisible to prestimulus analyses. As noted above, we previously identified that an intervening phase reset causes problems for relating pre-stimulus phase to neural processing occurring after the reset occurs^28^, as is likely the case for all but the lowest levels of visual processing. This problem arises because prestimulus phase analyses assume a statistical correspondence between phase in the prestimulus period and phase at the time of task-related neural processing, and a phase reset breaks this correspondence. It has also been shown that the onset of an event-related potential can corrupt phase estimates at nearby times, producing blind-spots in phase-behaviour analyses^72^.

Another limitation on the ability for phase studies to provide meaningful information about the role of phase in cognitive processing is suggested by the fact that our significant phase effects occurred roughly 500ms prior to stimulus onset. How could the phase of oscillations have an influence on processing before the stimulus appeared, but not during the intervening period until target onset? Correspondence between prestimulus phase and behavioural outcomes, as shown by our significant POS analysis, implies that the oscillations continued from the time of our effects until the stimulus was processed. We did not observe a relationship between the oscillation and behaviour at these times, despite the result at earlier times telling us the relationship was present. Our analysis of oscillatory power over this same period suggests this was most likely due to low oscillatory power reducing our ability to reliably estimate phase over this period^65–67^. When amplitude is zero, phase values become random. Well before amplitude approaches zero, however, in the presence of aperiodic noise, low signal-to-noise ratio of the data reduces the accuracy of phase estimates ^65,67^. Low signal-to-noise ratio may be particularly problematic for studies of oscillation phase in areas of cognition such as spatial attention and temporal expectation, as these phenomena are known to produce suppression of alpha oscillations ^79–81,86,95,96^, and thus are likely to produce less reliable phase estimates.

In summary, our results provide a clear indication that neural oscillation phase influences interactions between distinct neural populations in visual areas. These results provide a significant advance on prior detection studies, showing an influence of neural oscillations on feature-specific decision processes, as would be expected to exist if neural oscillations play a fundamental role in mediating neural communication and computation.

## Methods

### Participant Details

Forty healthy adults (26 self-identified females, 14 self-identified males, mean age = 23.69 years, SD = 3.86 years, all right-handed) participated in this study. The age data of four participants were lost due to a technical error. One participant was excluded from analysis due to problems with the EEG recording resulting in large artefacts that could not be corrected using standard procedures. Three additional participants were excluded because the behavioural titration procedure failed to converge. No experimental data was collected for these participants. There were 36 participants (24 self-identified females) in the final analysed dataset. Sex differences were not examined in this study as there are no sex differences in the perception of the tilt illusion^98^, and our study would not have been sufficiently powered for such a comparison. Participants were recruited through The University of Queensland’s research participation platform, gave written informed consent before participating, and were compensated at a rate of $20 per hour for their time. All experimental procedures were approved by the Human Research Ethics Committee at The University of Queensland (2021/HE002284).

### Method Details

#### Stimuli and Apparatus

All stimuli were generated using MATLAB (MathWorks, R2020B) with the Psychophysics Toolbox^99,100^ (version 3.0.17) and presented on a gamma-corrected LCD monitor (VIEWPixx 3D, VPixx Technologies; 1920 x 1080 pixels, 22.5-inch, 120 Hz). A viewing distance of 57 cm was maintained using a chin- and forehead-rest. A black cardboard annulus with a central hole of approximately 26° was placed in front of the monitor, such that the stimuli were visible through the annulus, but the participants were not able to view any portion of the monitor frame. A black cloth was draped over surrounding equipment to remove all external cues to vertical. Central fixation was controlled using a video-based infra-red eye tracker (EyeLink 1000 Plus; SR Research). The eye tracker was calibrated before each task block using the standard five-point calibration. Participants responded to the stimulus using a USB keyboard.

On each trial, a fixation target comprising an inner black dot (0.17°; RGB: 0, 0, 0) and outer white ring (0.3°; RGB: 255, 255, 255) appeared for 1500–2000ms on a grey background (RGB: 160, 160, 160; Figure 1). Next, a circular central test grating (5° diameter) and a surrounding annular grating (20° diameter; absent in the ‘surround-absent’ titration condition, see below) were presented simultaneously for 8.33ms. Extremely brief presentations were employed because longer stimuli would overlap many different oscillatory phase angles. Past research has shown that the tilt illusion is robust when stimuli are presented briefly^35^. Both central and surround gratings had a spatial frequency of 1cpd and Michelson contrast of 0.6. Prior to the main task, we titrated the central grating to each participant’s subjective vertical angle for our two main conditions (surround ±30°) and also included a surround-absent condition to quantify participants’ bias in the perception of vertical when the illusion was absent (further titration details below). During the main task, the surround grating was oriented at ±30°, and the central grating was presented at each individual’s titrated vertical angle for the given surround orientation. A thin grey ring (0.3° width) between the central grating and the surrounding annulus was presented to reduce perceptual interference at the point where the centre and surround gratings met. Participants had an untimed response period in which to indicate whether they perceived the central grating to be tilted leftward or rightward relative to vertical, using their right hand on the left and right arrows of the keyboard. They did not receive feedback on their responses.

After the response was made for a given trial, a 200ms circular masking stimulus was presented to prevent visual aftereffects from one trial influencing perception on the next. The masking stimulus comprised a combination of six gratings, angled every 30° from 30° to 180°, with each grating at a different random phase on every trial. It is important to note, this was not a visual mask in the classical sense, in that it did not serve to reduce the visibility of the target. Stimuli in this task were clearly visible, the response period was untimed, and the mask was presented after the response was made.

#### Titration

Participants completed the experiment in a dark, electro-magnetically shielded room. Following EEG setup (see EEG Acquisition), participants received written and oral instructions. They then completed two stages of titration to determine the threshold angle at which they were equally likely to report the central grating as rotated left and right of vertical (their ‘subjective vertical’) for each of three conditions: +30° surround (right-surround condition), -30° surround (left-surround condition), and a central grating with no surround grating (no-surround condition).

In the first stage of titration, each stimulus was titrated separately in two 1-up-1-down staircases. This provided a total of six staircases which were interleaved in their presentation. For each condition, both staircases began with a central orientation of ±15° which was adjusted in 1° increments after each response. All staircases ended after 10 reversals. The threshold for each staircase was calculated as the average orientation of the final 6 reversals. This value was averaged for the two staircases of each condition.

To ensure accurate estimates of participants’ subjective vertical thresholds (as phase-opposition analyses [see below] require a similar number of trials in each condition^36^, with the conditions of interest here being leftward versus rightward responses), a second stage of titration was conducted. This titration again used two staircases for each condition, with each staircase beginning at the threshold orientation derived from the first titration ±0.5°, then adjusted in 0.125° increments after each response. These staircases continued for 15 reversals and the final threshold estimate for each staircase was again the average orientation of the final 6 reversals. The final threshold estimate for each condition was the average threshold of its two staircases. The final thresholds in the left- and right-surround conditions were used as the central orientations in the main task (Figure 1B and **Table S1**).

#### Main Task

The main task proceeded in identical fashion to the titration task, with EEG recorded during each trial. To account for changes in illusion magnitude due to fatigue or habituation, if participants made the same response to one condition five times in a row, the angle of the central stimulus was adjusted 0.125° in the opposite direction. This adjustment was performed to ensure that similar numbers of leftward and rightward responses were made in each condition, as required for phase opposition analysis. The main task consisted of 10 experimental blocks of 80 trials each, for a total of 800 trials (excluding titration trials). Self-paced break periods were provided after each block.

### Quantification and Statistical Analysis

#### Behavioural Analyses

Magnitude of the tilt illusion was quantified by subtracting the threshold central orientation for the surround-absent condition from the left- and right-surround conditions. This allowed us to compare the illusion magnitudes between the two conditions, correcting for any bias in participants’ perceptions of vertical. When quantifying change in illusion magnitude over the course of the experiment, we compared the raw, unadjusted thresholds for the left- and right-surround conditions to those achieved at the end of the experiment, because there was no surround-absent condition presented in the main experiment to allow for subtraction. For all EEG analyses, leftward and rightward response data from the two surround conditions were combined by recoding responses in each condition as illusion-consistent (away from the flanker direction, i.e., rightward responses on left-surround trials, and leftward responses on right-surround trials) or illusion-inconsistent (rightward responses on right-surround trials, and leftward responses on left-surround trials). The justification for this is that the tilt illusion is a repulsive illusion^29^, and thus a neural state that produced an increase in the magnitude of the illusion would be expected to result in more responses away from the surround direction for a given central stimulus, whereas a state that reduced the magnitude of the illusion would result in more responses toward the surround direction.

#### EEG Recording

An Active Two system (BioSemi) recorded continuous EEG data, digitised at 1024 Hz with 24-bit A/D conversion. Sixty-four active Ag/AgCl electrodes covered the scalp, placed according to the standard international 10-10 system^101^ using a nylon cap. The Common Mode Sense and Driven Right Leg electrodes served as the reference and ground during recording. Eye muscle activity was recorded using bipolar horizontal electro-oculographic (EOG) electrodes at the outer canthi of each eye and bipolar vertical EOG electrodes above and below the left eye. Bilateral mastoid electrodes served as import references.

#### EEG Preprocessing

EEG pre-processing was performed offline using the EEGLAB Toolbox^102^ for MATLAB, and analyses were performed using custom MATLAB scripts, the specparam toolbox^48^, and code from VanRullen^36^. Data were down-sampled to 256 Hz, high-pass filtered with EEGLAB’s default FIR filter at 0.1 Hz and re-referenced to the average of all electrodes. Bad electrodes were detected by EEGLAB’s default kurtosis-based detection algorithm (threshold: 5 *SD*s) and removed for later interpolation. Eye blinks which overlapped stimulus presentation were identified as any trial containing >100 µV deviation on any ocular electrode during -200 to +200ms relative to stimulus onset. These trials were removed from subsequent analyses (*M* = 25.02 trials, *SD* = 35.85, range = 0–159). Data were re-referenced to the average of all electrodes and epoched from -2000 to +1000ms relative to stimulus onset. The data were baseline corrected by subtracting the mean voltage from -100 to 0ms from all timepoints. Using infomax ICA, data were then examined for artefacts of non-neural origin, including eye blinks, eye movements, muscle activity, line noise, etc. Components were evaluated by visual inspection with the assistance of the SASICA plugin for EEGLAB^103^, which incorporates the ADJUST^104^ and FASTER^105^ plugins. On average, 15.75 components were removed per participant (SD = 5.06, range = 5–32). Following ICA cleaning, channels which had previously been removed from the data were interpolated with spherical spline interpolation from surrounding channels. An average of 4.08 electrodes were interpolated per participant (SD = 1.79, range = 1–8).

#### ROI Selection

The tilt illusion is well understood to result from local inhibitory interactions in early visual processing^30–32^. As such, for all EEG analyses we focussed on a region of interest (ROI) concentrated on posterior central electrodes (electrodes Iz, Oz, O1, O2, and POz). The ROI was chosen apriori but conformed well to the topography of the early stimulus evoked response (**Figure S1.**) For analyses at the ROI, signals from the ROI channels were averaged prior to spectral decomposition. To allow for the possibility of effects outside our ROI location, we also report the results of analyses performed at each individual electrode, corrected for multiple comparisons by controlling the False Discovery Rate^61^ (FDR).

#### Power Spectrum Parameterisation

EEG data were separated into periodic and aperiodic components using the specparam toolbox version 1.0.0^48^. Specparam iteratively fits the EEG power spectrum with a 1/f^x^ function, representing the aperiodic component of the data, and some number of gaussians above that function, which capture the oscillatory bumps above the aperiodic spectrum. To extract single-trial and trial-averaged power spectra, we applied the Welch method to Hann tapered data from the pre-stimulus period (-600 to 0ms) to decompose single-trial data into frequencies between 0 and 40 Hz in increments of 0.25 Hz. The specparam procedure was run on frequency data from 3-40 Hz with the following settings: peak width limits: 0.5–12 Hz; maximum number of peaks: 3; minimum peak height: 0 (which sets a peak height threshold relative to the scale of the data); peak threshold: 2; aperiodic mode: ‘fixed’. Fits were quite good, with average R^2^ values of 0.94 (SD = .03). For alpha power analyses, power was determined by taking the magnitude parameter for peaks in the alpha range (8–14 Hz). Trials in which no peak in the alpha range was detected were assigned an alpha power of 0. To avoid including participants with little-to-no alpha in the alpha power analyses, we also calculated the trial-averaged spectra for every channel for every participant. Participants for whom no alpha was detected at more than half of their channels (3 participants), or who had no alpha detected at any ROI channel (1 participant), were excluded from alpha power analyses.

#### Logistic Mixed Effects of Alpha Power and Aperiodic Parameters

Alpha power, spectral slope and offset measures from single trials were standardised within participants by converting to z-scores and were entered as predictors into a logistic mixed effects analysis with a random effect of participants, predicting responses (illusion-consistent vs illusion-inconsistent) on single trials.

#### Phase Analysis

For the phase analyses, trial data from -2000 to +1000ms relative to stimulus onset were decomposed into 38 integer frequencies from 3 to 40 Hz. This decomposition was performed via convolution in the frequency domain with complex Morlet wavelets with a width of 3 cycles at 3 Hz, increasing linearly to 7 cycles at 40 Hz. Data were then re-epoched to the pre-stimulus period from -600 to 0ms. Phase at each frequency and timepoint was calculated as the angle of the complex frequency representation resulting from wavelet analysis. Phase opposition analysis (see below) was performed at all times (-600 to 0 ms) and frequencies for signals from the ROI (averaged, then wavelet decomposed). An exploratory analysis was also performed at all frequencies, times, and electrodes, corrected for multiple comparisons with FDR (**Figure S3**).

Phase analyses were performed using the phase opposition sum (POS) metric^36^. This metric compares the sum of the inter-trial phase consistency (ITPC) for two separate trial groups— here, illusion-consistent versus illusion-inconsistent responses— against the ITPC for both groups collapsed together, using the following formula:

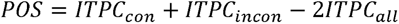

Where *ITPCcon* is the ITPC of illusion-consistent trials, *ITPCincon* is ITPC of illusion-inconsistent trials, and *ITPCall* is ITPC of all trials combined. ITPC is calculated as:

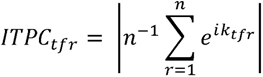

Where *e^ik^* indicates the complex polar representation of phase values from each time, *t*, frequency, *f,* and trial, *r*. This formula computes the absolute value of the vector-average of a series of unit-length vectors of differing phase angles, *k*.

When trial groups cluster at different phases of an oscillation (e.g., at a peak vs. a trough), POS yields a positive score. If phase angles are random or are clustered at the same angle in both conditions, POS will approach zero. POS values were converted to p-values using a non-parametric permutation test with 1000 permutations. The mean and standard deviation of the permuted null distribution were characterised to calculate a z-score for the true POS value and this z-score was converted to a p-value by reference to the cumulative normal distribution. Individual participant p-values were combined to produce a group-level p-value at each frequency and timepoint using Stouffer’s method^106^. The group-level p-values were adjusted for multiple comparisons using FDR correction^61^.

To quantify the effects of phase on behaviour we employed an unbiased phase alignment procedure^15,17,20^. The phase values from each time and frequency within significant POS clusters were divided into seven bins of equal width from -π to π radians, and the proportion of illusion-consistent responses was calculated for the trials in each bin. Next, the bin containing the lowest proportion of illusion consistent responses was assigned to the central bin at 0 radians. The remaining phase bins were then rotated to maintain their position relative to the bin assigned to 0 radians. The central bin that had been used for phase alignment was then excluded from statistical analysis, as it was selected based on its proportion of illusion-consistent responses and was therefore biased. The remaining bins, which were aligned only by virtue of their relationship to the central bin, contain no bias. If a phase effect is present, then a monotonic increase in illusion-consistent responses should emerge across bins with increasing distance from the central bin. If a phase effect is absent, however, then the bin selected for alignment should have had the lowest proportion of illusion-consistent responses by chance, and there should be no systematic pattern of responses across the remaining bins after alignment.

After excluding the central bin, six bins remained. We averaged the behavioural data of the two bins closest to the central bin to produce a single value for the ‘near-central’ phases. We also averaged the data from the most distant bins to produce a single value for the ‘far-from-central’ phases. A one-tailed repeated measures *t*-test was used to compare the proportion of illusion-consistent responses between the near-central and far-from-central phases, averaged across each significant time and frequency identified in the POS analysis. This test was one-tailed because the expected direction of phase effects is determined by the phase-alignment procedure. Note: the significance of any phase effects is already determined by the POS analysis prior to any alignment analysis. The purpose of the alignment is to visualise the relationship between phase and behaviour, rather than to provide an independent test.

## Acknowledgements

The authors would like to acknowledge the following funding sources: WJH was supported by the Australian Research Council (DE190100136). JBM was supported by the National Health and Medical Research Council (GNT2010141). AMH was supported by the Australian Research Council (DE220101019).

