## Supplementary Materials for "Effects of oscillation phase on discrimination performance in a visual tilt illusion"

**Table S1. Uncorrected subjective vertical thresholds (degrees)**

| Participant | CCW Surround | No Surround | CW Surround |
| --- | --- | --- | --- |
| 2 | -7.30 | -1.56 | 3.81 |
| 3 | -7.45 | 2.25 | 9.99 |
| 4 | -10.32 | -3.33 | 0.49 |
| 5 | -6.39 | 0.79 | 4.01 |
| 6 | -7.62 | -1.66 | 1.09 |
| 7 | -2.41 | 2.34 | 7.07 |
| 9 | -8.29 | -2.14 | 4.10 |
| 10 | -8.85 | -2.00 | -1.46 |
| 11 | -3.75 | 2.55 | 11.92 |
| 12 | -7.41 | 1.44 | 10.51 |
| 13 | -5.21 | -0.79 | 4.48 |
| 14 | -4.67 | 0.97 | 3.44 |
| 15 | -12.83 | -3.56 | 1.98 |
| 17 | -4.97 | 1.62 | 5.38 |
| 18 | -8.65 | -3.36 | 0.66 |
| 19 | -5.91 | -0.91 | 3.17 |
| 20 | -6.04 | -0.61 | 3.57 |
| 21 | -8.67 | 1.28 | 4.24 |
| 22 | -7.90 | -2.11 | 0.18 |
| 23 | -11.66 | -2.68 | 5.21 |
| 24 | -5.15 | 0.63 | 4.71 |
| 25 | -4.65 | 1.50 | 7.38 |
| 26 | -7.77 | 0.40 | 8.40 |
| 27 | -7.81 | -2.08 | 1.14 |
| 29 | -4.31 | 1.36 | 4.40 |
| 30 | -2.99 | 4.24 | 13.94 |
| 31 | -6.65 | -1.78 | 2.18 |
| 32 | -4.88 | -2.21 | 2.25 |
| 33 | -2.78 | -0.09 | 2.42 |
| 34 | -2.31 | 0.80 | 4.99 |
| 35 | -7.79 | -1.12 | 4.53 |
| 36 | -10.24 | -3.66 | 2.82 |
| 37 | -4.26 | 0.13 | 4.72 |
| 38 | -9.55 | -3.46 | 1.39 |
| 39 | -6.60 | -1.75 | 3.80 |
| 40 | -3.88 | -1.35 | 3.43 |
| Mean (SD): | -6.61 (2.58) | -0.55 (2.02) | 4.34 (3.34) |

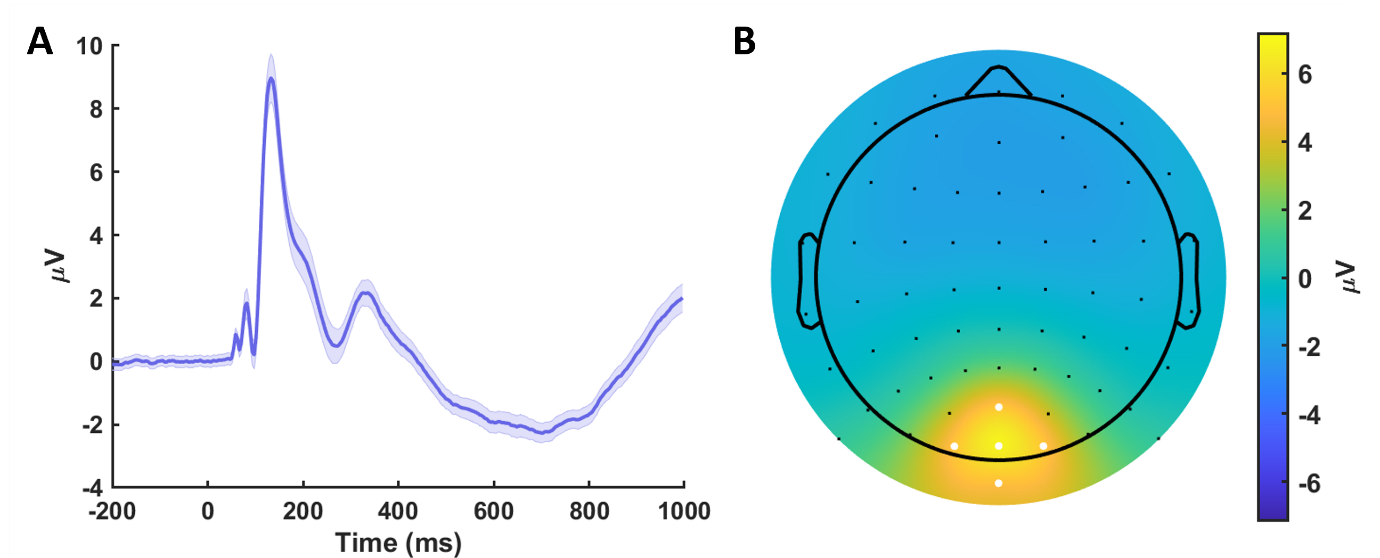

**Figure S1. Event related potential.** A – Event related potential (ERP) averaged across the ROI and across conditions. Shading represents within-participants SEM. B – Topography of the ERP averaged between 100 and 200ms. White dots indicate ROI electrodes.

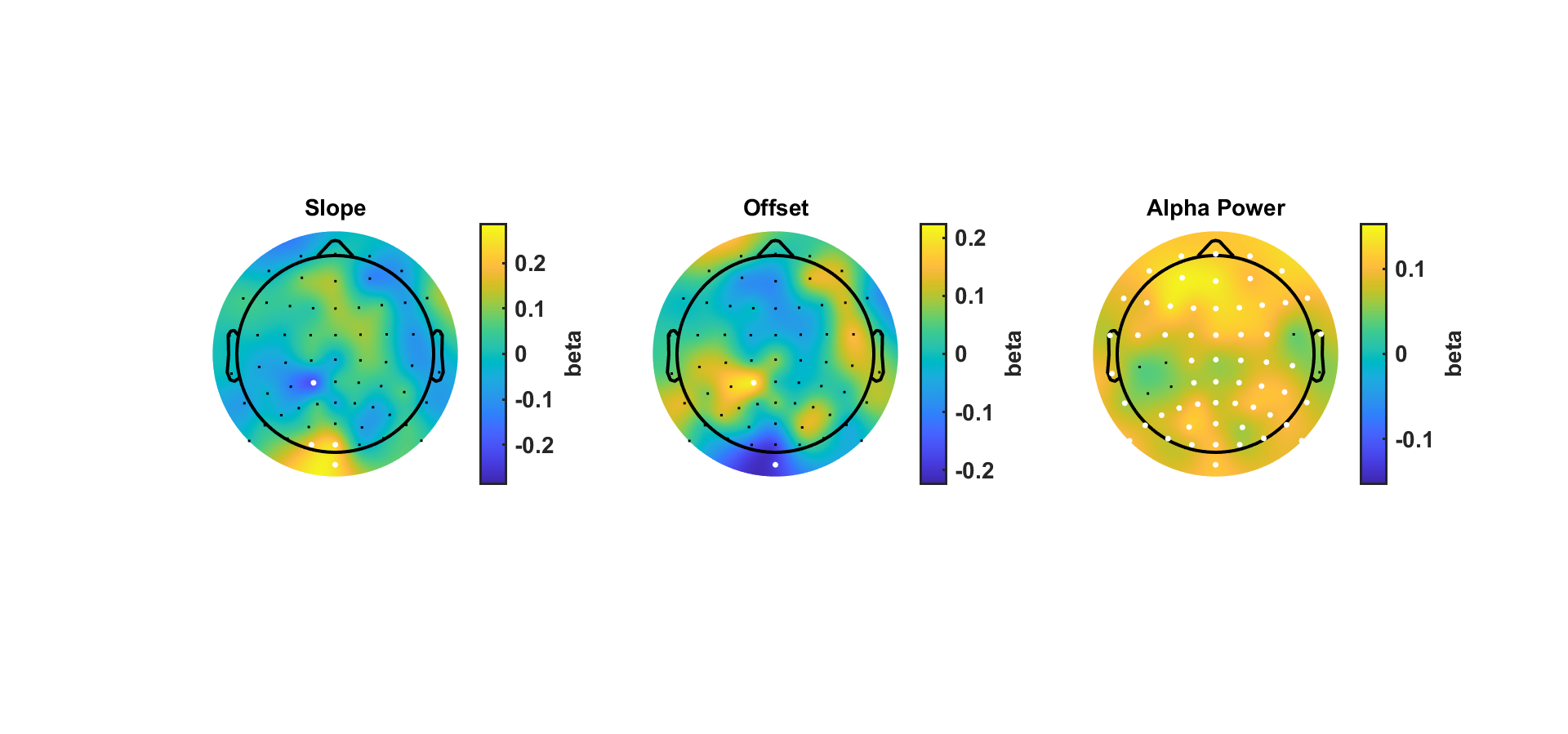

**Figure S2. Logistic mixed effects at each channel.** For each of the three parameters of interest we performed an exploratory logistic mixed effects analysis at each channel. White dots indicate significant electrodes after FDR correction. Slope: 4 significant electrodes. Offset: 2 significant electrodes. Alpha power: 58 significant electrodes. Note, each topography has a different colour scale.

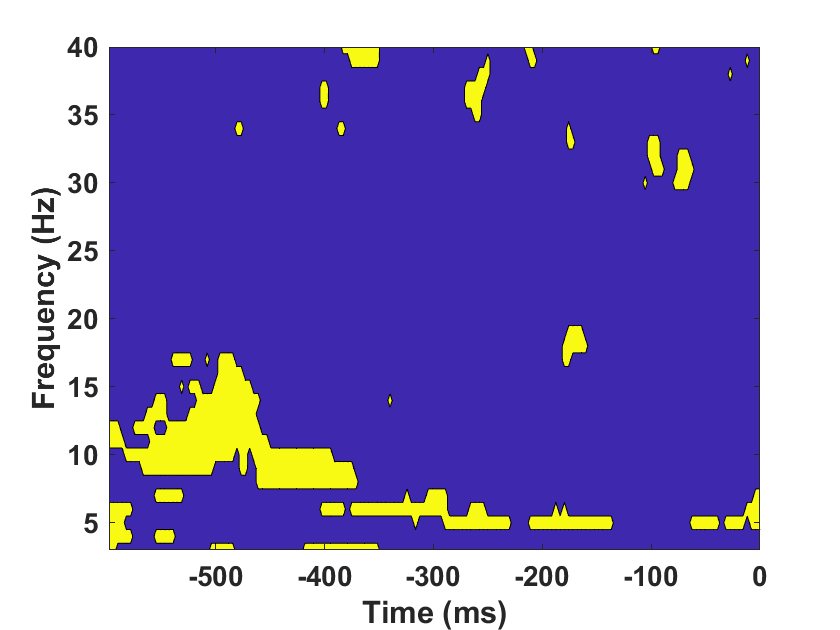

**Figure S3. POS across all electrodes.** Yellow points indicate times/frequencies with significant POS at one or more channels. *P*-values were corrected with FDR across all times, frequencies, and channels.
